# Rapid protection of nonhuman primates against Marburg virus disease using a single low-dose VSV-based vaccine

**DOI:** 10.1101/2022.09.17.508396

**Authors:** Kyle L. O’Donnell, Friederike Feldmann, Benjamin Kaza, Chad S. Clancy, Patrick W. Hanley, Paige Fletcher, Andrea Marzi

## Abstract

Marburg virus (MARV) is the causative agent of Marburg virus disease (MVD) which has a case fatality rate up to ~90% in humans. Recently, there were cases reported in Guinea and Ghana highlighting this virus as a high-consequence pathogen potentially threatening global public health. There are no licensed treatments or vaccines available today.

We used a vesicular stomatitis virus (VSV)-based vaccine expressing the MARV-Angola glycoprotein (VSV-MARV) as the viral antigen. Previously, a single dose of 1×10^7^ plaque-forming units (PFU) administered 7 days before challenge resulted in uniform protection from disease in cynomolgus macaques. Here, we sought to lower the vaccination dose to allow for more doses per vial in an emergency outbreak situation. We administered 1×10^5^ or 1×10^3^ PFU 14 days before challenge and achieved uniform protection in both groups. When we administered 1×10^3^ PFU 7 days before challenge, vaccination resulted in uniform protection with no detectable viremia. Antigen-specific IgG responses were induced by both vaccine concentrations and were sustained until the study endpoint. Neutralizing antibody responses and antibody-dependent cellular phagocytosis were observed with both vaccination doses and timelines. The cellular response after vaccination was characterized by early induction of NK cell activation. Additionally, antigen-specific memory T cell subsets were detected in all vaccination cohorts indicating that while the primary protective mechanism of VSV-MARV is the humoral response, a functional cellular response is also induced.

Overall, this data highlights VSV-MARV as a viable and fast-acting MARV vaccine candidate suitable for deployment in emergency outbreak situations and supports its clinical development.

**One Sentence Summary:** A single low dose of VSV-MARV administered 14 or 7 days before challenge protects NHPs uniformly from lethal disease.

## INTRODUCTION

Marburg virus (MARV) is a member of the *Filoviridae*, the same family as Ebola virus (EBOV), which has a 19 kb (-) single-stranded RNA genome encoding 7 proteins. The mature viral particles are filamentous in structure and exit the cell through budding from the surface (*1*). MARV first emerged in 1967 in Marburg, Germany, and Belgrade, former Yugoslavia and from then on it has caused sporadic outbreaks in parts of Africa (*2, 3*). The largest outbreak occurred in 2004/05 in Angola in which 252 cases were identified and 227 fatalities recorded (*4*). The 2021-22 outbreaks in Western Africa, specifically Guinea and Ghana, emphasized the risk of introducing this high-consequence pathogen into a new geographical area (*5, 6*). Indeed, computational modeling estimates that 105 million people are at risk of MARV infection in Africa and Madagascar (*7*). The clinical manifestation of Marburg virus disease (MVD) progresses from initial non-specific flu-like symptoms to petechiae, delirium, multi-organ dysfunction, and massive hemorrhaging. There is neither an approved vaccine nor treatment for MVD, and due to the highly pathogenic nature and effective human-to-human transmission via bodily fluids, MARV is on the list of priority pathogens by the World Health Organization (*8*).

Vesicular stomatitis virus (VSV)-based vaccines have shown promising pre-clinical success against several viral families including *Coronaviridae, Arenaviridae, Paramyxoviridae*, and *Filoviridae* (*9*). The greatest success to date of the platform is the EBOV vaccine approved by the US Food and Drug Administration (FDA) and the European Medicines Agency (EMA) under the name “Ervebo” (also known as VSV-EBOV and rVSV-ZEBOV)(*10, 11*). Similar to VSV-EBOV, a vaccine expressing the MARV glycoprotein (GP) as the primary viral antigen (VSV-MARV) has demonstrated uniform protection in nonhuman primates (NHPs). As previously demonstrated, a single dose of 1×10^7^ plaque-forming units (PFU) uniformly protected NHPs when lethal challenge occurred 28, 14, or 7 days post-vaccination (DPV) (*12–14*). This vaccine has also demonstrated the ability to protect NHPs in a post-exposure therapeutic challenge setting (*15*). Due to the potential for a large outbreak as indicated by computational modeling, and the reality that only limited GMP vaccine doses are available for clinical application, we sought to determine if reducing the dose from 1×10^7^ PFU to 1×10^5^ PFU or 1×10^3^ PFU would retain the high protective efficacy in a 14 or 7 DPV challenge setting, respectively. We demonstrate that NHPs that received a single dose of VSV-MARV at 1×10^5^ PFU or 1×10^3^ PFU were uniformly protected from lethal challenge 14 DPV. When the low dose of 1×10^3^ PFU was administered there was also uniform protection from lethal disease with challenge 7 DPV. All control animals succumbed to infection with hallmarks of MVD.

## RESULTS

### Single low-dose VSV-MARV Vaccination Protects NHPs Within 7 Days from Lethal Disease

We sought to assess the minimum dose that would maintain the protective efficacy of VSV-MARV previously demonstrated. Groups of 4 NHPs were vaccinated with 1×10^5^ PFU or 1×10^3^ PFU VSV-MARV intramuscularly (IM) and challenged with a lethal IM dose of 1,000 PFU MARV Angola 14 DPV. Vaccination resulted in complete protection from severe disease. A control vaccine, VSV-EBOV, at 1×10^5^ PFU afforded no protection and NHPs succumbed to disease 6 or 7 days post-challenge (DPC) (Fig. 1A). Only the control NHPs developed signs of MVD reflected in the increased clinical scores; other parameters we evaluated revealed elevated liver enzyme levels, and high titer viremia in control NHPs (Fig. 1B-E). Only the control NHPs developed cytokine levels suggestive of the characteristic MVD-associated cytokine storm (fig. S1). Next, we shortened the time between vaccination and challenge as previously described (*14*) to assess if efficacious immunity at a low dose can be achieved within one week. NHPs were challenged IM 7 DPV with 10^3^ PFU of VSV-MARV which resulted in 100% survival and no signs of MVD (Fig. 2A). In contrast, control NHPs vaccinated with 10^3^ PFU of VSV-EBOV developed signs of MVD (Fig. 2B-E) and a cytokine storm (fig. S1). They reached euthanasia criteria 6 and 7 DPC.

**Figure 1.**
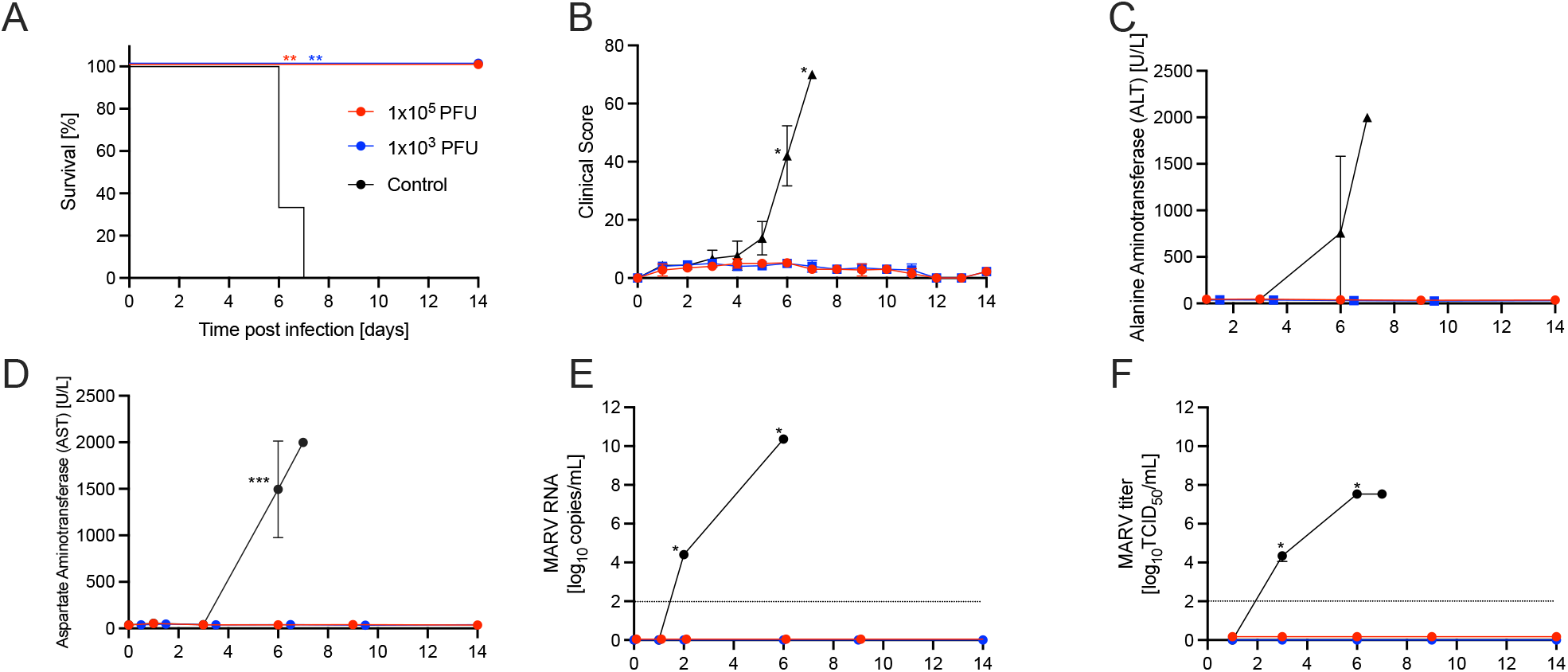
VSV-MARV protects NHPs within 14 days from lethal challenge. NHPs were vaccinated with a single intramuscular dose of VSV-MARV or VSV-EBOV. (A) Survival and (B) clinical scores during the acute disease phase are shown. Levels of (C) alanine aminotransferase (ALT) and (D) aspartate aminotransferase (AST) in the serum of MARV-infected NHPs. MARV viremia assessed (E) by RT-qPCR and (F) viral titration. Geometric mean and geometric SD are depicted in E, F. Statistical significance of survival was determined by Mantel-Cox test, other data were evaluated by Mann-Whitney test. Statistical significance is indicated as *p* < 0.001 (***), *p* < 0.01 (**), and *p* < 0.05 (*). Lines indicate limit of detection.

**Figure 2.**
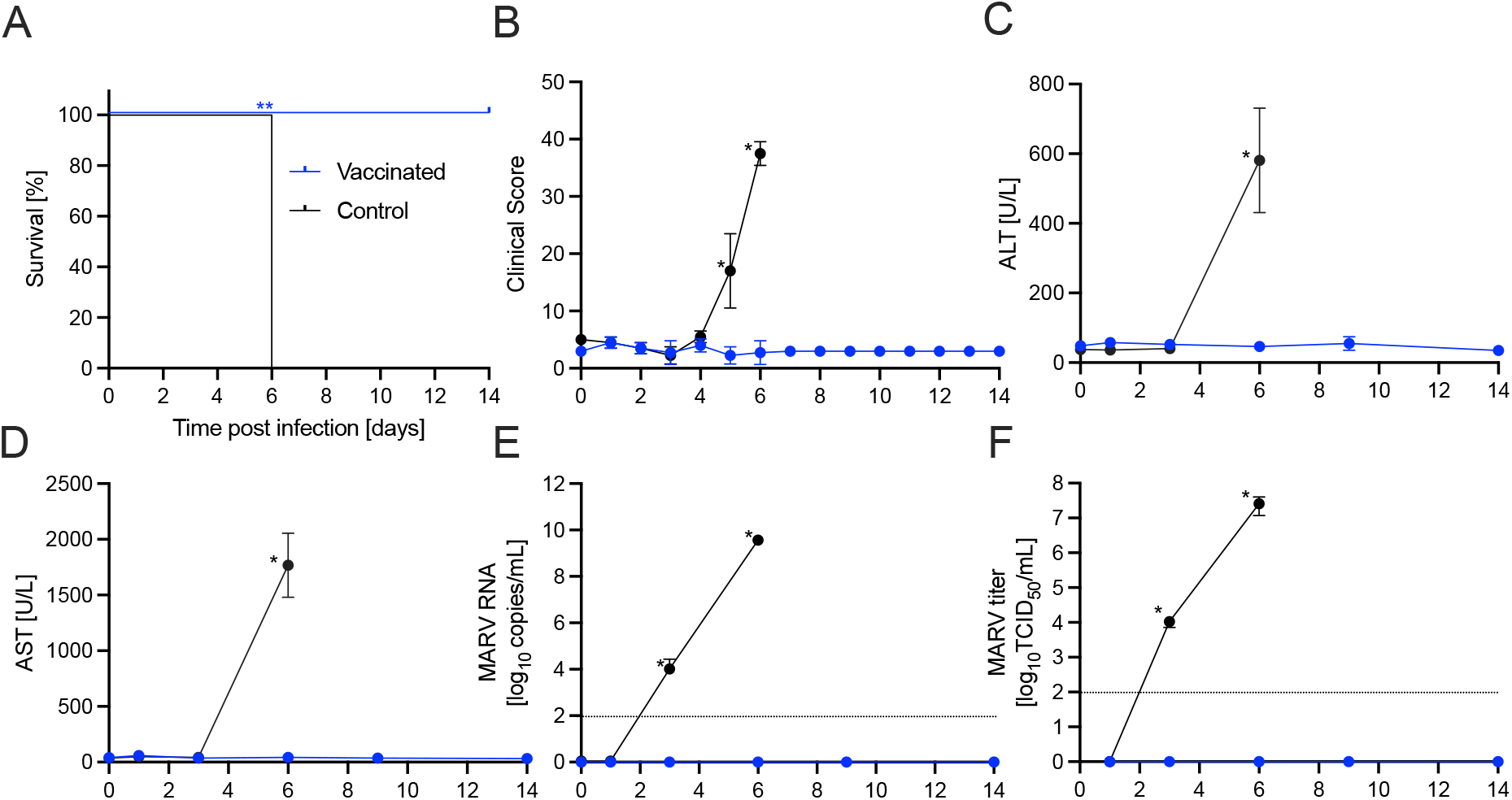
VSV-MARV protects NHPs within 7 days from lethal challenge. NHPs were vaccinated with a single intramuscular dose of VSV-MARV or VSV-EBOV. (A) Survival and (B) clinical scores during the acute disease phase are shown. Levels of (C) alanine aminotransferase (ALT) and (D) aspartate aminotransferase (AST) in the serum of MARV-infected NHPs. MARV viremia assessed (E) by RT-qPCR and (F) viral titration. Geometric mean and geometric SD are depicted in E, F. Statistical significance of survival was determined by Mantel-Cox test, other data were evaluated by Mann-Whitney test. Statistical significance is indicated as *p* < 0.01 (**), and *p* < 0.05 (*). Lines indicate limit of detection.

### VSV-MARV-Vaccinated NHPs Develop Antigen-Specific and Multifunctional Humoral Responses

It has previously been established that the primary mediation of protection with VSV-based vaccines is antibody-driven (*16, 17*). Therefore, we sought to determine the antigen specificity of both the IgM and IgG responses. Serum samples collected throughout the studies determined that the MARV GP-specific IgM response for all doses peaked around 14 DPV (Fig. 3A, 4A). The MARV GP-specific IgG response showed varying kinetics depending on vaccination dose and timing. NHPs vaccinated with 1×10^5^ PFU responded with a rapid IgG increase for 14 days followed by a sustained response until the study endpoint (Fig. 3B). NHPs vaccinated with 1×10^3^ PFU and challenged 14 DPV showed a similar rapid increase as the 1×10^5^ PFU group, however, at 6 DPC there is a significant drop in GP-specific IgG which rebounded quickly (Fig. 3B). NHPs challenged 7 DPV with 1×10^3^ PFU did not show an increase in MARV GP-specific IgG until 6 DPC, after which there was a steady increase until the study endpoint (Fig. 4B). We also measured the IgG response to another MARV antigen not part of the vaccine, the virion protein 40 (VP40). Despite the lack of viremia in any of the VSV-MARV vaccinated groups, the NHPs developed low levels of VP40-specific IgG at the study end point indicating challenge virus exposure (Fig. 3C, 4C). As expected, titers were higher in NHPs that received the lower vaccine dose.

**Figure 3.**
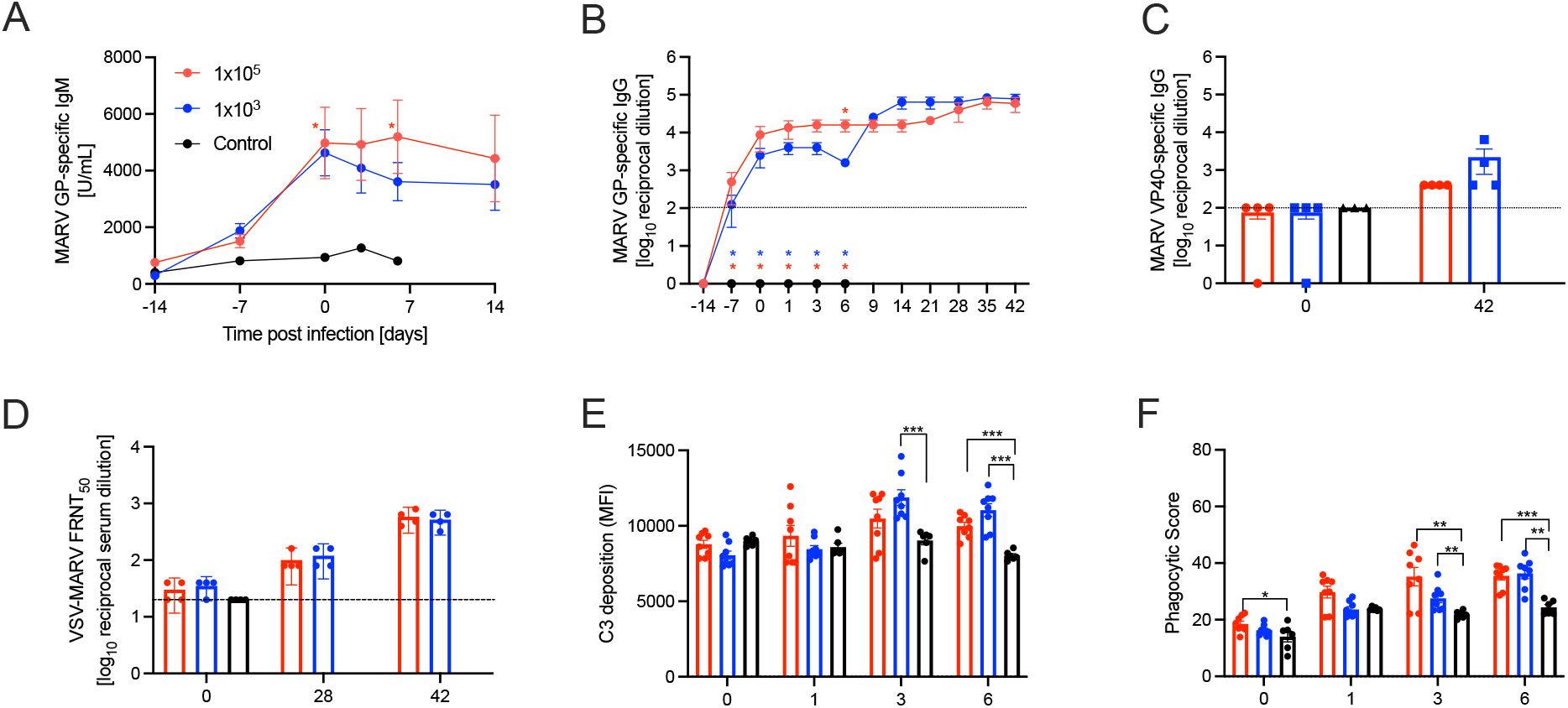
Humoral response and antibody functions in NHPs vaccinated 14 days before challenge. Concentrations of circulating of MARV GP-specific (A) IgM or (B) IgG, and (C) MARV VP40-specific IgG. Functionality of the antigen-specific responses assessed by (D) neutralization (median fluorescence reduction neutralization titer, FRNT50), (E) antibody-dependent complement deposition (ADCD), and (F) antibody-dependent cellular phagocytosis (ADCP). Mean and SEM are depicted. Statistical significance as determined by the Mann–Whitney test is indicated as *p* < 0.001 (***), *p* < 0.01 (**), and *p* < 0.05 (*). Lines indicate limit of detection.

**Figure 4.**
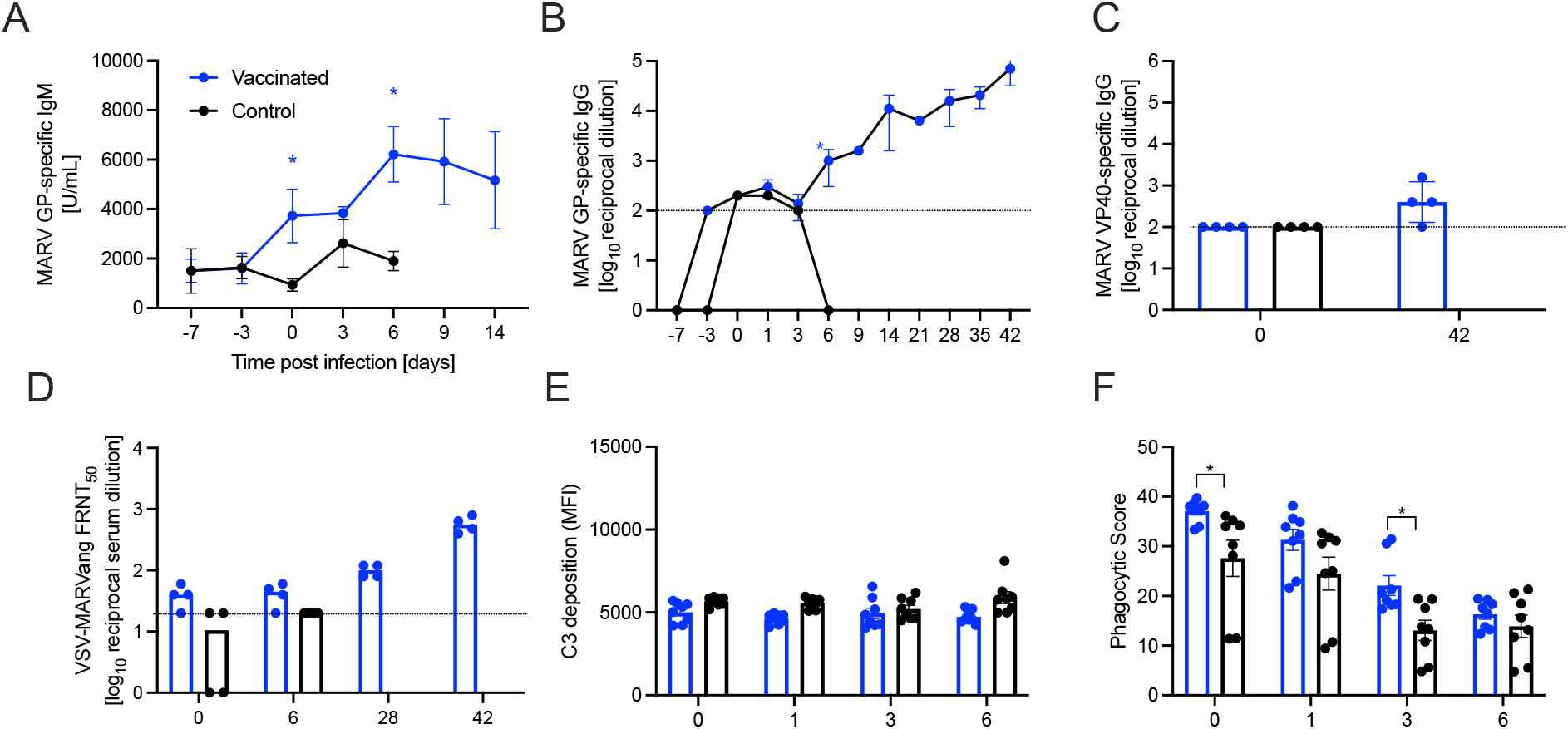
Humoral response and functionality in NHPs vaccinated 7 days before challenge. Concentrations of circulating of MARV GP-specific (A) IgM or (B) IgG, and (C) MARV VP40-specific IgG. Functional capabilities of the antigen-specific responses assessed by (D) neutralization (FRNT50), (E) antibody-dependent complement deposition (ADCD), and (F) antibody-dependent cellular phagocytosis (ADCP). Mean and SEM are depicted. Statistical significance as determined by the Mann–Whitney test is indicated as *p* < 0.05 (*). Lines indicate limit of detection.

The functionality of the humoral response was characterized with a neutralization assay using replication-competent VSV-MARV expressing GFP (VSV-MARV-GFP). We determined that there was no significant difference in neutralization titers between vaccinated and control NHPs regardless of vaccine dose or timing of vaccination at the time of challenge. However, and in line with the MARV GP-specific IgG, vaccinated NHPs showed an increase in viral neutralization titers over time until the study endpoint (Fig. 3D, 4D). Additionally, antibody Fc effector functions were assessed including antibody-dependent complement deposition (ADCD) and antibody-dependent cellular phagocytosis (ADCP). NHPs vaccinated with 1×10^5^ PFU demonstrated a significantly higher ADCP activity at 3 DPC as well as ADCD activity 6 DPC, respectively. The serum of NHPs vaccinated with 1×10^3^ PFU 14 days before challenge demonstrated a significantly higher ADCD and ADCP activity at 3 DPC (Fig. 3E, F). When the low dose of 1×10^3^ PFU was administered 7 days before challenge, ADCD was not different compared to control, and only transiently more ADCP activity was measured on 0 and 3 DPC (Fig. 4E-F).

### The timing of the VSV-MARV Vaccination Alters the NK Cell Response in NHPs

Next, we investigated the innate immune response with a focus on the NK cell response as a potential contributor to the rapid protection. Cryo-preserved PBMCs isolated from whole blood samples collected on 0, 14 and 28 DPC enabled us to characterize the functional phenotypes of the NK cell compartment throughout the study. In addition to antigenic stimulation (MARV GP-specific peptide pool), we also assessed if the humoral response played a role in activating NK cells by antibody-dependent cellular cytotoxicity (ADCC). NHPs vaccinated with 1×10^5^ PFU VSV-MARV and challenged 14 DPV responded with a rapid increase of CD107a and Granzyme B in the general CD16+ PBMC NK cell compartment at 0 DPC only (Fig. 5A-C). In contrast, NHPs vaccinated with 1×10^3^ PFU regardless of challenge time point did not demonstrate any significant increases in NK cell functionality compared to the control cohorts (fig. S2). We then assessed the ability of the humoral response to induce ADCC in each vaccination group at the time of challenge. NHPs vaccinated with 1×10^5^ PFU VSV-MARV did not demonstrate any significant increase in ADCC activity (Fig. 5D-F). NHPs vaccinated with 1×10^3^ PFU and challenged 14 DPV responded with an increase in CD107a and Granzyme B in the CD8+ NK cell compartment at the time of challenge only (Fig. 5D-F). This phenotype however was not apparent in the NHPs vaccinated with 1×10^3^ PFU and challenged 7 DPV. A longitudinal comparison of the activated NK cells response in this cohort overtime suggests that optimal NK cell activation occurred 14 DPC (21 DPV) (fig. S2).

**Figure 5.**
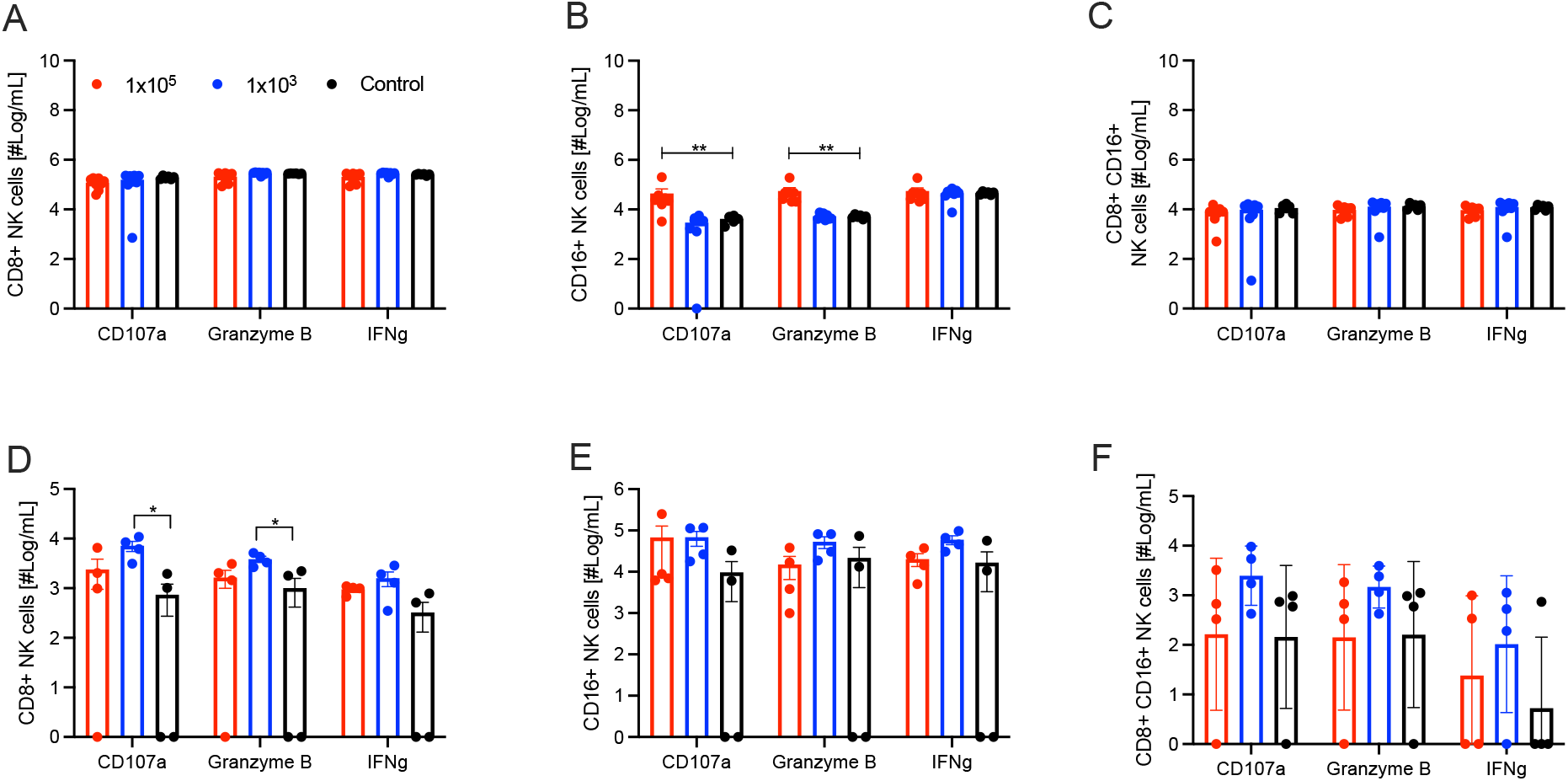
NK cell responses 14 days after VSV-MARV vaccination. Number of circulating MARV GP-specific activated NK cells on 0 DPC in the cohort challenged 14 DPV. (A) CD8^+^CD3^-NK cells^ expressing CD107a, Granzyme B, and IFNγ. (B) CD16^+^CD3^-^ NK cells expressing CD107a, Granzyme B, and IFNγ. (C) CD8^+^ CD16^+^ CD3^-^ NK cells expressing CD107a, Granzyme B, and IFNγ. Functional capacities of the antigen-specific MARV GP IgG to induce antibody dependent cellular cytotoxicity (ADCC). (D) CD8^+^CD3^-^ NK cells expressing CD107a, Granzyme B, and IFNγ. (E) CD16^+^CD3^-^ NK cells expressing CD107a, Granzyme B, and IFNγ. (F) CD8^+^ CD16^+^ CD3^-^ NK cells expressing CD107a, Granzyme B, and IFNγ. Mean and SEM are depicted. Statistical significance as determined by the Mann–Whitney test is indicated as *p* < 0.01 (**), and *p* < 0.05 (*).

### VSV-MARV Vaccination Induces an Activated CD4+ T cell Bias

As stated previously, it is well-established that VSV-based vaccines mediate protection primarily via the humoral response and cellular responses play a limited role. The potential of memory antiviral T cell formation and the extent to which T cells facilitate the humoral response was our next focus of analysis. The CD4+ T cells demonstrated a vaccine-dependent activation response with higher amounts of naïve CD4+ T cells expressing IFNγ for both vaccination doses in this cohort. NHPs vaccinated with 1×10^3^ PFU demonstrated higher amounts of activated EM with IFNγ+ and TNFα+ (Fig. 6A-D). The increased activation phenotype also included EM-RE CD4 T cells with higher amounts of IFNγ and TNFα accumulation in the 1×10^3^ PFU vaccination cohort (Fig. 6A-D). Both vaccine doses resulted in sustained CD4+ T cell functionality with a slight difference only in the EM activation state at 14 DPC in NHPs vaccinated with 1×10^5^ PFU; these NHPs presented with a significantly higher amount of CD69+ EM CD4+ T cells at that time (Fig. 6E-G). NHPs vaccinated and challenged 7 DPV did not demonstrate any significant changes in any of the T cell compartments analyzed on 0 DPC when compared to control (fig. S3). However, in both the CD4+ and CD8+ T cells the maturation of the cellular response in the vaccinated animals is apparent for the naïve, CM, and EM-RE compartments as there is a drop in all populations of activated cells on 14 DPC and then a recovery by 28 DPC. Surprisingly, we did not observe any activated EM cells in this cohort (fig. S4).

**Figure 6.**
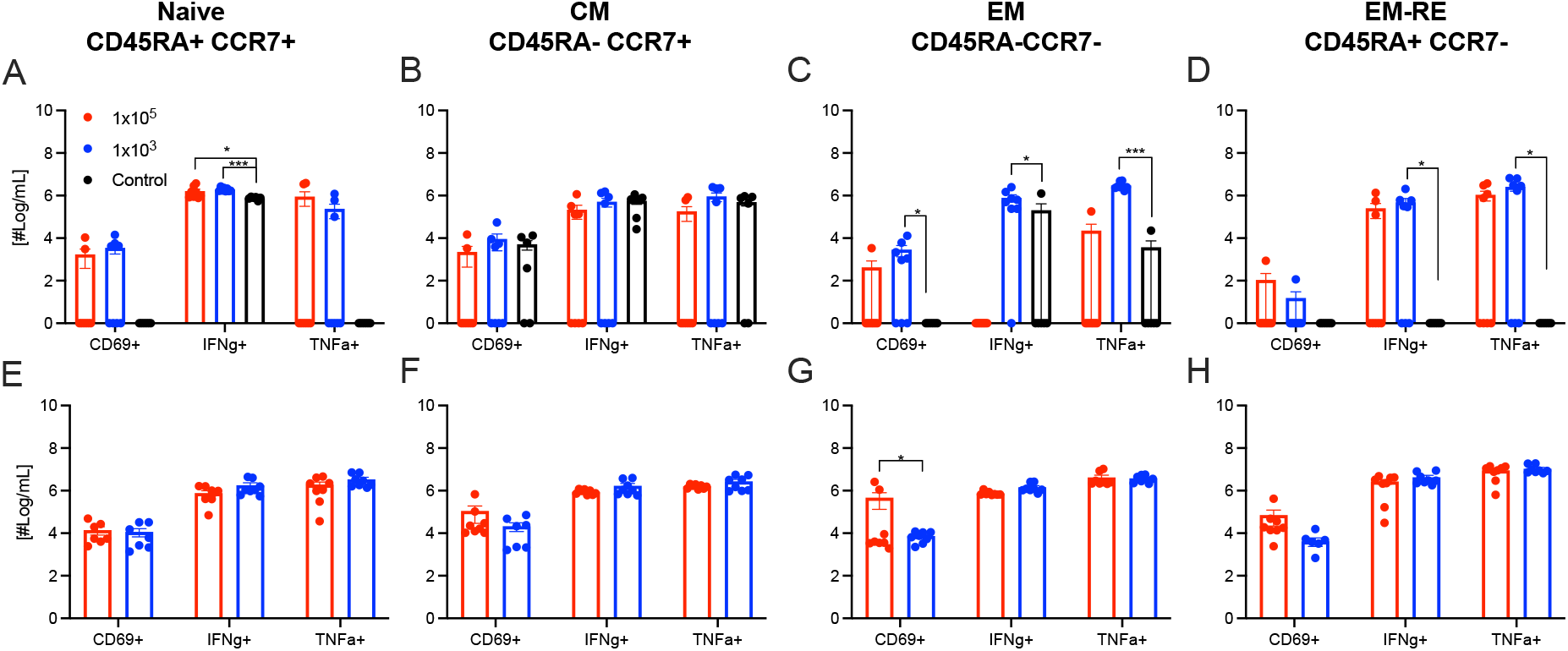
CD4^+^ T cell responses in NHPs vaccinated 14 days before MARV challenge. Number of circulating MARV GP-specific activated T cells on 0 DPC (A-D) and 14 DPC (E-H). (A) Naïve, (B) Central memory (CM), (C) Effector memory (EM), (D) Effector memory re-expressing CD4 T cells expressing CD69, IFNγ, and TNFα on 0 DPC. (E) Naïve, (F) CM, (G) EM, or (H) Effector memory re-expressing CD4 T cells expressing CD69, IFNγ, and TNFα on 14 DPC. Mean and SEM are depicted. Statistical significance as determined by the Mann–Whitney test is indicated as *p* < 0.001 (***) and *p* < 0.05 (*).

We determined that in the CD8+ T cell compartment at the time of challenge (0 DPC), none of the vaccination regimens induced a strongly activated CD8+ T cell population (fig S4). In contrast, the control cohort showed higher amounts of cellular cytokine levels in the central memory (CM), effector memory (EM), and effector memory re-expressing (EM-RE) populations (fig. S4). There was a significant difference between the NHPs vaccinated with 1×10^5^ PFU or 1×10^3^ PFU and challenged 14 DPV in the EM-RE IFNγ+ and TNFα+ cells at 14 DPC. A similar trend was observed at 28 DPC in the EM populations when NHPs vaccinated with 1×10^3^ PFU presented with higher amounts of IFNγ+ and TNFα+ cells (fig. S4).

## DISCUSSION

In a previous MARV study, we established that a single dose of VSV-MARV at 1×10^7^ PFU uniformly protect NHPs within 7 days (*14*). In this study, we assessed whether a single low-dose of VSV-MARV could still rapidly protect vaccinated NHPs from MVD. We determined that a single dose as low as 1×10^3^ PFU uniformly protected NHPs when challenged 7 DPV. Our results add to the growing body of evidence that this platform provides rapid protection and is suitable for deployment during an outbreak. A dose-down study using a VSV-based EBOV vaccine administered IM 28 days before challenge demonstrated 100% efficacy with as little as 10 PFU in NHPs (*18*). We have demonstrated that the VSV-MARV remain fast-acting even at a lower dose highlighting the potential to extend the number of doses from one vial of the limited GMP-manufactured supply (*14, 18*). Based on the pre-clinical data presented here, and the dose-down study with the VSV-EBOV (*18*), we believe a dose-down phase one clinical trial would be warranted to determine the dose necessary for immunogenicity. While VSV-MARV and VSV-EBOV are very similar in mediating protection, there are also differences in the kinetic of induction of immunity and the ideal vaccine doses might be different for each vaccine. In addition, compilation of a complete antibody functionality profile would allow for a better understanding of the protective capabilities, not relying solely on neutralizing which is known to not correlate with protection for this vaccine platform (*19, 20*). VSV-based vaccines are live-attenuated vaccines that have in general caused mild adverse effects in humans such as irritation, headache, fatigue, fever, chills, myalgia, and arthralgia (*21*). Reducing the administered vaccine dose while retaining protective efficacy would reduce adverse events and their severity thereby optimizing not only the doses available per vial, but the experience of the vaccinees as well.

Protection using VSV-based vaccines against filovirus infections is primarily mediated by the antigen-specific humoral response (*16, 22*). The MARV GP-specific IgM and IgG responses peaked 14 DPV, which coincided with MARV challenge in our first study. These findings are similar to what has previously been reported for VSV-MARV and VSV-EBOV NHP studies (*14, 23*). When challenge occurred only 7 DPV with the low dose, the MARV GP-specific IgG response did not show the typical plateau 14 DPV, rather it continued to increase steadily for the duration of the study. It is unclear at this point which aspect of the MARV challenge stimulates this response. Analysis of the antibody functionality of VSV-MARV vaccine studies has been limited to neutralization assays which do not correlate with disease outcomes after MARV infection (*19*). Indeed, we assessed the neutralization capacity of the humoral response here and found similar trends with no significant difference between the VSV-MARV vaccinated groups and the control NHPs at the time of challenge. However, the neutralizing response matures as the study progresses as demonstrated by the increase in neutralizing titers at study end.

Our further investigation of Fc effector functions included complement activation, cellular phagocytosis, and cellular cytotoxicity. Data from a clinical cohort of Ebola virus disease (EVD) survivors demonstrated the induction of ADCD regardless of the neutralization titer suggesting protective benefits of ADCD against filovirus infections (*24*). This study determined that the low dose VSV-MARV vaccination 14 days before challenge resulted in significantly higher complement activation levels 17 DPV. However, the primary ADCD response occurred 20 DPV for both doses. When the low dose was administered only 7 days before challenge, we did not see an increase in complement deposition, suggesting that antibody class switching, and maturation may be required to fully activate the complement cascade via ADCD.

The next Fc effector function we assessed was the induction of cellular phagocytosis (ADCP) which directly reflects the activation of the innate immune system. Innate myeloid cells are the primary phagocytic cells, and upon phagocytosis become activated while degrading the engulfed particle. The cohort challenged 14 DPV demonstrated increased phagocytosis after challenge closer to peak disease of the control animals. The 7 DPV cohort had an inverse response showing a decrease of phagocytosis as the study progressed. This observation could be due to antibody consumption to control the infection with the low-dose vaccination and challenge 7 DPV. Indeed, MARV GP-specific IgM levels remained constant for the first few days after challenge and MARV GP-specific IgG titers were not above control levels until 6 DPC supporting this hypothesis. Similar results were found in the assessment of therapeutic monoclonal antibodies demonstrating the importance of phagocytic functions (*25, 26*).

The final functional response we investigated was the induction of ADCC. We found that only the low-dose vaccinated NHPs challenged 14 DPV demonstrated an induction of a cytotoxic phenotype of CD8^+^ CD3^-^ NK cells expressing increased amounts of degranulation marker CD107a and Granzyme B accumulation. Surprisingly, there was no indication of ADCC in the vaccinated NHPs challenged 7 DPV with no activation phenotypes detected in the CD16^+^ CD3^-^ or CD8^+^ CD16^+^CD3^-^ compartments. A previous study with VSV-MARV in NHPs included transcriptomic analysis indicating that the activation level of the innate immune responses and Fc receptor-mediated signaling at the time of challenge may play a role in protection (*14*). Antibody class switching is essential for Fc effector functions; therefore, it is not unexpected that vaccinated NHPs challenged 7 DPV had a lower functional response and a more immature antibody repertoire. However, our analysis demonstrates that several antibody Fc effector functions contribute to protection. These findings are consistent with previous reports from EVD survivors, in which polyfunctional humoral responses were demonstrated in survivors and, albeit to a lesser extent, in patients who succumbed to disease (*20, 27*). These findings are not only present in clinical samples and vaccine studies, but the development of monoclonal antibody therapies have also heavily researched polyfunctional responses and have demonstrated that neutralization alone does not lead to optimal protection *in vivo.* Increased efforts to design monoclonal antibody therapies with polyfunctional capabilities have generated the most efficacious treatments (*26–31*).

Previous studies have highlighted the importance of the innate immune response in the rapid protection conferred by VSV-based vaccines (*14, 23, 32, 33*). We sought to investigate the role of NK cells regarding innate immunity activation using flow cytometry-based assays. We analyzed PBMCs and found a significantly activated NK cell response in the CD16^+^ CD3^-^ compartment with significantly higher amounts of CD107a and Granzyme B on 0 DPC for the high-dose vaccinated NHPs challenged 14 DPV compared to controls. There was no difference between the vaccine groups on 14 and 28 DPC. This contrasts with the ADCC data which demonstrated activation of the low-dose group in the CD8^+^ CD3^-^ compartment, indicating that either the vaccine dose or the innate signature before challenge may skew the NK cell response. There was no difference in the NK cell response for vaccinated NHPs from the 7 DPV cohort at 0 DPC. Longitudinal analysis found that significantly higher NK cell function on 14 DPC compared to 0 and 28 DPC. The activated NK cell phenotype was associated with B cell help indicated by increased amounts of CD107a and IFNγ indicating that NK cell involvement post-vaccination requires a 14 day maturation period prior to challenge.

Although it has previously been shown that there is limited T cell contribution to protection against filoviruses (*14, 16*), we further characterized the cellular response performing T cell immune phenotyping. We sought to investigate the extent in which the T cells contributed to the maturation of the humoral response. We first investigated the cohort challenged 14 DPV and determined that both vaccine doses at 0 DPC (time of challenge) elicited significantly more naïve CD4^+^ expressing IFNγ than the control NHPs. The low-dose vaccinated NHPs had significantly more IFNγ and TNFα in the CD4^+^ EM and EM-RE populations at 0 DPC. This observation suggests that a lower vaccine dose may stimulate a more robust memory phenotype quicker after vaccination. The low-dose group continued to show increased CD4^+^ T cell activation compared to the high-dose group 14 and 28 DPC suggesting the low-dose elicited cellular response played a more significant role in protection. This difference could be attributed to the speed at which the protection occurred and virus was cleared resulting in a lack of antigen expression to further stimulate antigen-specific T cell expansion for the higher dose group. The same analysis was performed on PBMCs isolated from NHPs challenged 7 DPV. However, we did not detect any significant differences between the vaccinated and control groups suggesting that 7 days may not be enough time to stimulate the specific cellular response after vaccination sufficiently. Like the 14 DPV cohort, the speed at which the protection occurred may have limited the cellular response with the lack of antigen present to stimulate subset expansion. Collectively, and in line with previous research, we demonstrated that the T cell response is not the primary protective immunological component for VSV-based vaccination. Rather, the responses are supporting the humoral response, allowing for greater expansion and activation of plasma cells to bolster the antibody response.

This study had limitations including the lack of a decrease in protective efficacy with the low-dose vaccinated NHPs in the 7 DPV cohort. Future work will encompass the protective efficacy testing of even lower doses of VSV-MARV to determine the minimum dose necessary for protection within one week. This will facilitate an in-depth characterization of the innate cell responses which was focused on NK cell responses here. A broader innate cell phenotyping approach will give insight into other innate cells that contribute to the rapid protection conferred by VSV-based vaccines. All cellular phenotyping here was performed on cryopreserved PBMCs reflective of the cellular components that are circulating within the host at that specific point in time. We did not assess tissue-specific cellular responses which could play a pivotal role in early sites of viral replication. Finally, the sample size for each cohort was relatively low posing the risk of missing rare events and vaccine breakthrough.

Small outbreaks of MARV have increased in frequency as demonstrated by the recent cases in Guinea (2021) and Ghana (2022). In this populated area in West Africa, it may be only a matter of time before a large outbreak of MVD occurs as MARV has the potential to spread efficiently in areas with poor healthcare setting similar to EBOV which caused an epidemic in West Africa in 2013-2016 (*34*). While there is still no approved MARV vaccine available, the data presented here support for VSV-MARV to move into clinical development as soon as possible. The gathered data may support the use of this vaccine in an outbreak under emergency use authorizations.

## MATERIALS AND METHODS

### Ethics Statement

All the work involving infectious MARV was performed following standard operating procedures (SOPs) approved by the Rocky Mountain Laboratories (RML) Institutional Biosafety Committee (IBC) in the maximum containment laboratory at the RML, Division of Intramural Research, National Institute of Allergy and Infectious Diseases, National Institutes of Health. Procedures were conducted by trained personnel under the supervision of veterinary staff on animals anesthetized with ketamine. All efforts were made to ameliorate animal welfare and minimize animal suffering per the Weatherall report on the use of nonhuman primates in research (https://royalsociety.org/policy/publications/2006/weatherall-report/). Animal work was performed in strict accordance with the recommendations described in the Guide for the Care and Use of Laboratory Animals of the National Institute of Health, the Office of Animal Welfare, and the United States Department of Agriculture and was approved by the RML Animal Care and Use Committee (ACUC). Animals were housed in adjoining individual primate cages that enabled social interactions, under controlled conditions of humidity, temperature, and light (12-h light:12-h dark cycles). Food and water were available *ad libitum.* Animals were monitored and fed commercial monkey chow, treats, and fruit at least twice a day by trained personnel. Environmental enrichment consisted of commercial toys, music, and video. Endpoint criteria based on clinical score parameters as specified and approved by the RML ACUC were used to determine when animals were humanely euthanized.

### Animal Study Design

Nineteen male or female cynomolgus macaques (*Macaca fascicularis)* 3-5 years of age and 1.9-2.8 kg in weight were used for this study. Three groups of cynomolgus macaques (n=4 per vaccination group; n=3 for control) were vaccinated with a single IM injection of 1×10^5^ PFU or 1×10^3^ PFU VSV-MARV and challenged 14 DPV. Control NHPs were IM-vaccinated with 1×10^5^ PFU VSV-EBOV and challenged 14 DPV. Clinical exams including a blood draw on anesthetized NHPs were conducted on −14, −11, −7, 0, 1, 3, 6, 9, 14, 21, 28, 35 and 42 DPC. The second study entailed two groups of cynomolgus macaques (n=4 per group) which were vaccinated with 1×10^3^ PFU VSV-MARV or VSV-EBOV and challenged 7 DPV. Clinical exams including a blood draw on anesthetized NHPs were conducted on −7, −4, 0, 1, 3, 6, 9, 14, 21, 28, 35 and 42 DPC. An IM injection of 1,000 PFU MARV-Angola (confirmed by back-titration) as previously described (*13*) served as lethal challenge for all NHPs. The animals were observed at least twice daily for clinical signs of disease according to a RML ACUC-approved scoring sheet and humanely euthanized when they reached endpoint criteria. The study ended 42 DPC when all surviving animals were humanely euthanized.

### Cells and Virus

Vero E6 cells (*Mycoplasma* negative) were grown at 37°C and 5% CO2 in Dulbecco’s modified Eagle’s medium (DMEM) (Sigma-Aldrich, St. Louis, MO) containing 10% fetal bovine serum (FBS) (Wisent Inc., St. Bruno, Canada), 2 mM L-glutamine, 50 U/mL penicillin, and 50 mg/mL streptomycin (all supplements from Thermo Fisher Scientific, Waltham, MA). THP-1 (*Mycoplasma* negative) were grown at 37°C and 5% CO2 in Roswell Park Memorial Institute medium (RPMI) (Sigma-Aldrich, St. Louis, MO) containing 10% FBS (Wisent Inc., St. Bruno, Canada), 2 mM L-glutamine, 50 U/mL penicillin, and 50 mg/mL streptomycin (all supplements from Thermo Fisher Scientific).VSV vaccines and MARV-Angola were propagated in Vero E6 cells using DMEM supplemented with 2% FBS, L-glutamine, and penicillin/streptomycin. VSV-MARV was constructed in-house as previously described (*13*) and used for IM vaccination. VSV-MARV-GFP was constructed by adding the GFP gene between the MARV GP and VSV polymerase gene into the viral genome (*35*). VSV-MARV-GFP was recovered from this plasmid as described previously (*36*). MARV-Angola was obtained from the Public Health Agency of Canada (GenBank accession number KY047763 (*37*)), propagated on Vero E6 cells, titered, and stored in liquid nitrogen. All viruses were confirmed by sequencing.

### Hematology and Serum Chemistry

The total white blood cell, neutrophil, lymphocyte, and platelet counts were determined from EDTA blood with the IDEXX ProCyte DX analyzer (IDEXX Laboratories, Westbrook, ME) serum biochemistry including aspartate aminotransferase (AST), alkaline phosphatase (ALP), alanine aminotransferase (ALT), glucose, creatinine, and total bilirubin was analyzed on a Vetscan 2 using Preventive care profile disks (Abaxis, Union City, CA).

### Viral Load Quantification

Blood samples were extracted using the QIAamp Viral RNA Mini Kit (Qiagen, Hilden, Germany) according to manufacturer specifications. Tissues, a maximum of 30 mg each, were processed and extracted using the RNeasy Mini Kit (Qiagen) according to manufacturer specifications. One step RT-qPCR for genomic viral RNA was performed using specific primer-probe sets to MARV L gene, and the QuantiFast Probe RT-PCR +ROX Vial Kit (Qiagen), in the Rotor-Gene Q (Qiagen). Forward primer CCTTGCCTTCCGATATGAATTT. Reverse primer TCACACCATAACATCGATTACAGTAGTC. Probe 6FAM-CGCGGCATTTCA-BBQ. Five μL of each RNA extract were run alongside dilutions of MARV standards with a known concentration of RNA copies. Concentrations were determined utilizing the Q-Rex 1.1.04 software with the absolute quantification plugin.

### MARV titers

Viremia was determined from EDTA whole blood samples using Vero E6 cells (*Mycoplasma* negative). Cells were seeded in 48-well plates the day before titration. On the day of titration, blood samples were thawed, and 10-fold serial dilutions were prepared in DMEM without supplements. Media was removed from cells and inoculated in triplicate with each dilution. After one hour, DMEM supplemented with 2% FBS, penicillin/streptomycin, and L-glutamine was added, and cells were incubated at 37°C. Cells were monitored for cytopathic effect (CPE) and a 50% tissue culture infectious dose (TCID50) was calculated for each sample employing the Reed and Muench method (*38*).

### Antigen-Specific Humoral Responses

Post-challenge NHP sera were inactivated by gamma-irradiation (4 MRad) (*39*) and removed from the maximum containment laboratory according to RML SOPs approved by the RML IBC. The MARV GP-specific IgM titers in serum samples were determined at 1:250 dilution using ELISA kits following manufacturer’s instructions (Alpha Diagnostics, San Antonio, TX). The MARV GP and VP40 IgG ELISAs were developed in-house. MARV-Angola GPdTM was obtained from IBT Bioservices (Gaithersburg, MD) and MARV VP40 was purified from transfected 293T cell supernatant. Nunc Maxisorp Immuno plates (Thermo Fisher Scientific) were coated with 50 μl of 1 μg/mL of antigen in PBS o/n and ELISAs were performed as described previously (*14*). The optical density (OD) at 405 nm was measured using a GloMax® explorer (Promega). The OD values were normalized to the baseline samples obtained with naïve NHP serum and the cutoff value was set as the mean OD plus three times the standard deviation of the blank.

### Quantification of antibody effector functions

Assays for antibody effector functions were adapted from previously established protocols (*40*). Post-challenge NHP sera were inactivated by gamma-irradiation (4 MRad) (*39*) and removed from the maximum containment laboratory according to RML SOPs approved by the RML IBC. Recombinant MARV-Angola GPdTM (IBT Bioservices) was tethered to Fluospheres NutrAvidin-Microspheres yellow-green or red (Thermo Fisher Scientific, Waltham, MA) using the EZ-link Micro Sulfo-NHS-LC-Biotinylation kit (Thermo Fisher Scientific).

#### ADCD

Serum samples were heat-inactivated at 56°C for 30 min then diluted in DMEM and applied to the conjugated beads for one hour at 37°C. After, guinea pig complement (Cedarlane, Burlington, Canada) was added for 30 minutes. The bead complexes were washed with FACS buffer and stained with anti-C3c-FITC. Data were acquired on a FACS Symphony (BD, Franklin Lakes, NJ) and analyzed in FlowJo v10.

#### ADCP

Serum samples were heat-inactivated at 56°C for 30 min then diluted in DMEM and applied to the conjugated beads for one hour at 37°C. The serum bead mixture was then transferred to a plate of THP-1 cells for 1 hour at 37°C. Data were acquired on a FACS Symphony (BD) and analyzed in FlowJo v10.

#### ADCC

Nunc Maxisorp Immuno plates (Thermo Fisher Scientific) were coated with 1 μg/mL recombinant soluble MARV-Angola GPdTM (Alpha Diagnostics) in PBS (50 ul/well). Serum samples were heat-inactivated at 56°C for 30 min, diluted in DMEM and then mixed with the PBMCs isolated. The antibody-PBMC mixture was transferred to the coated plate and incubated for 24 hours at 37°C. The cells were then stained for NK cell immune responses utilizing Live/Dead-UV450, CD45-BV786, CD3-FITC, CD8-PeTexas Red, CD16-AF700, CD20-BV421, and CD107a-PE. Cells were fixed with 4% paraformaldehyde (PFA) and stained intracellularly with IFN-γ-PE-Cy7 and Granzyme B-APC diluted in Perm-Wash buffer (Biolegend). Sample acquisition was performed on a Cytoflex-S (Beckman Coulter, Brea, CA) and data analyzed in FlowJo V10.

### Neutralization

Post-challenge NHP sera were inactivated by gamma-irradiation (4 MRad) (*39*) and removed from the maximum containment laboratory according to RML SOPs approved by the RML IBC. The day before this assay, Vero E6 cells were seeded into 96 well plates. Serum samples were heat-inactivated at 56°C for 30 min and 5-fold serially diluted in DMEM. VSV-MARV-GFP was added in equal volumes at a MOI of 1, and the mixture was incubated for 1 hour at 37°C. The antibody-viral solution was then transferred to the cells and incubated for 24 hours at 37°C and 5% CO2. The cells were fixed with 4% PFA and resuspended in FACs buffer. Data were acquired on a FACS Symphony (BD) and analyzed in FlowJo v10.

### Serum Cytokine Quantification

Serum samples were diluted 1:2 in serum matrix for analysis using the Milliplex Non-Human Primate Magnetic Bead Panel as per the manufacturer’s instructions (Millipore, Burlington, MA). Concentrations for G-CSF, GM-CSF, IFN-γ, IL-1ra, IL-1β, IL-2, IL-4, IL-5, IL-6, IL-8, IL-10, IL-12/23 (p40), IL-13, IL-15, IL-17, IL-18, MCP-1, MIP-1α, MIP-1β, sCD40L, TGF-α, TNF-α, and VEGF were determined for all samples. Values below the limit of detection of the assay were assigned the value of 1.

### Cellular Phenotyping Assays

PBMCs were isolated from whole blood samples using Histopaque®-1077 (Sigma-Aldrich) and separated according to manufacturers’ instructions. Isolated PBMCs were resuspended in FBS with 10% DMSO and frozen at −80°C until analysis.

For T cell response analysis, cells in duplicate were stimulated with 2μg/ml MARV GP peptide pool, media, cell stimulation cocktail (containing PMA-Ionomycin, Biolegend), or SARS-CoV-2 nucleocapsid peptide pool together with 5μg/ml Brefeldin A (Biolegend) for 16 hours. Following, cells were surface stained with Live/Dead-UV450, CD45-BV786, CD3-FITC, CD4-PerCP Cy5.5, CD8-PeTexas Red, CD69-AF700, CCR7-BV605, and CD45-RA-APC. Cells were fixed with 4% PFA and stained intracellularly with IFN-γ-BV421 and TNFα-PE diluted in Perm-Wash buffer (Biolegend).

NK cell immune responses were measured following the same method, but cells were surface stained with Live/Dead-UV450, CD45-BV786, CD3-FITC, CD8-PeTexas Red, CD16-AF700, CD20-BV421, and CD107a-PE. Cells were fixed with 4% PFA and stained intracellularly with IFN-γ-PE-Cy7 and Granzyme B-APC diluted in Perm-Wash buffer (Biolegend). Sample acquisition was performed on a Cytoflex-S (Beckman Coulter) and data analyzed in FlowJo V10.

## List of Supplementary Materials

fig. S1. **Cytokine responses in control and vaccinated NHPs after MARV challenge.**

fig. S2. NK cell responses after MARV challenge.

fig. S3. T cell responses after MARV infection in NHPs vaccinated 7 days before challenge.

fig. S4. CD8^+^ T cell responses after MARV infection in NHPs vaccinated 14 days before challenge.

## Acknowledgments

We thank the staff of the Rocky Mountain Veterinary Branch (NIAID) for their support of the NHP study. We also thank the staff supporting the maximum containment laboratory operation at RML.

## Funding

This work was supported by the Intramural Research Program NIAID, NIH.

## Author contributions

Conceptualization: KLO, AM

Methodology: KLO, PF

Investigation: KLO, PF, BK, FF, CSC, PWH, AM

Visualization: KLO

Funding acquisition: AM

Project administration: AM

Supervision: AM

Writing – original draft: KLO

Writing – review & editing: KLO, AM

## Competing interests

The authors declare that the research was conducted in the absence of any commercial or financial relationships that could be construed as a potential conflict of interest.

## Data and materials availability

All additional data are available in the main text or the supplementary materials.

---

**Figure S1.**
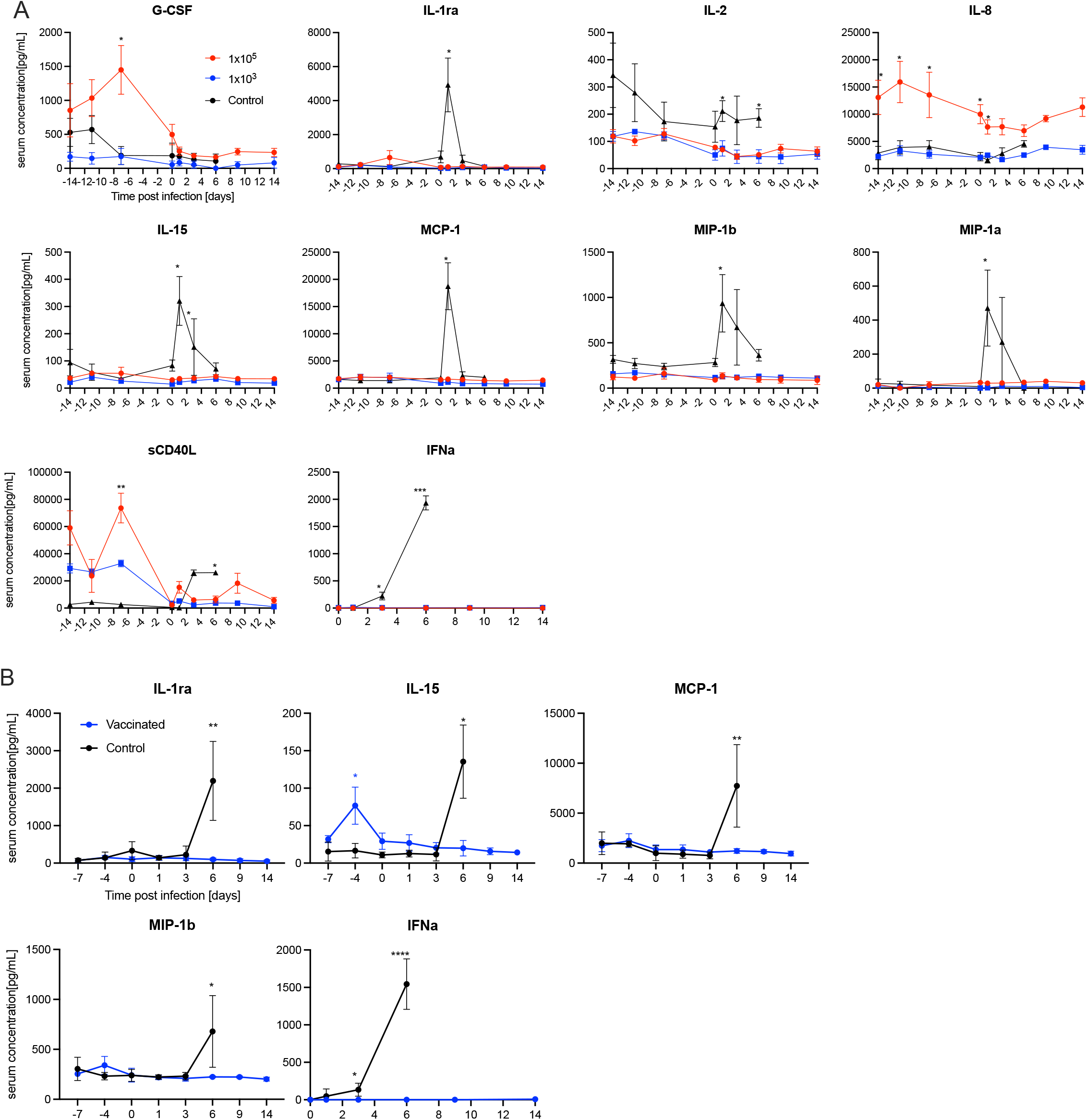
Cytokine responses in control and vaccinated NHPs after MARV challenge. The concentration of select cytokines in the serum of vaccinated and control NHPs over time. (**A**) NHPs challenged 14 DPV; (**B**) NHPs challenged 7 DPV. Mean and SEM are depicted. Statistical significance as determined by the Mann–Whitney test is indicated as *p* <0.0001 (****) *p* < 0.001 (***), *p* < 0.01 (**), and *p* < 0.05 (*).

**Figure S2.**
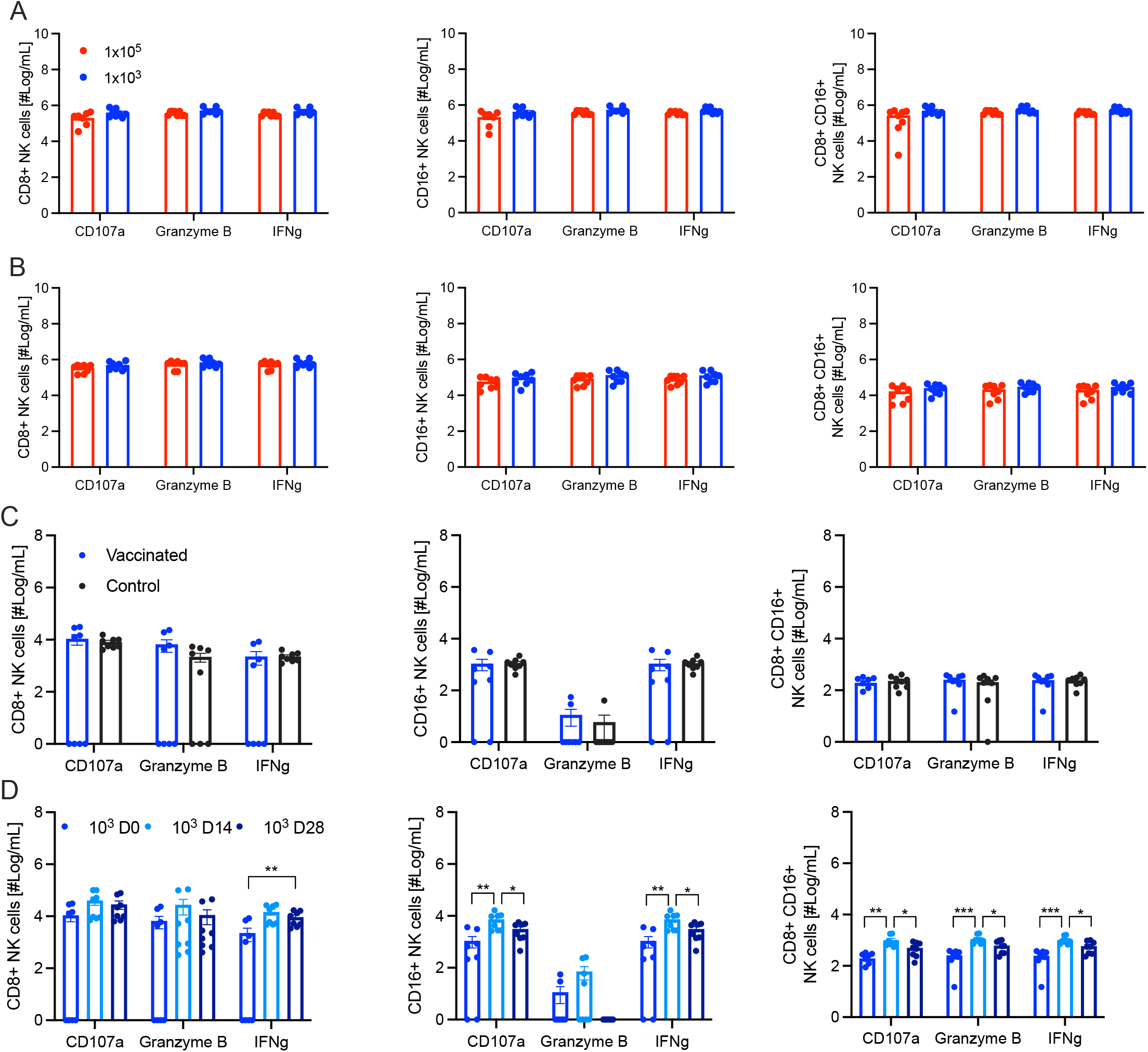
NK cell responses after MARV challenge. Number of circulating MARV GP-specific activated NK cells from the cohort challenged 14 DPV. NK cells at (**A**) 14 DPC or (**B**) 28 DPC expressing CD107a, Granzyme B, and IFNγ. Number of circulating MARV GP-specific activated NK cells from the cohort vaccinated 7 days before challenge. (**C**) 0 DPC comparing vaccinated and control NHPs. (**D**) Longitudinal comparison of the NK cells in the group challenged 7 DPV expressing CD107a, Granzyme B, and IFNγ. Mean and SEM are depicted. Statistical significance as determined by the Mann–Whitney test is indicated as *p* < 0.01 (**), and *p* < 0.05 (*).

**Figure S3.**
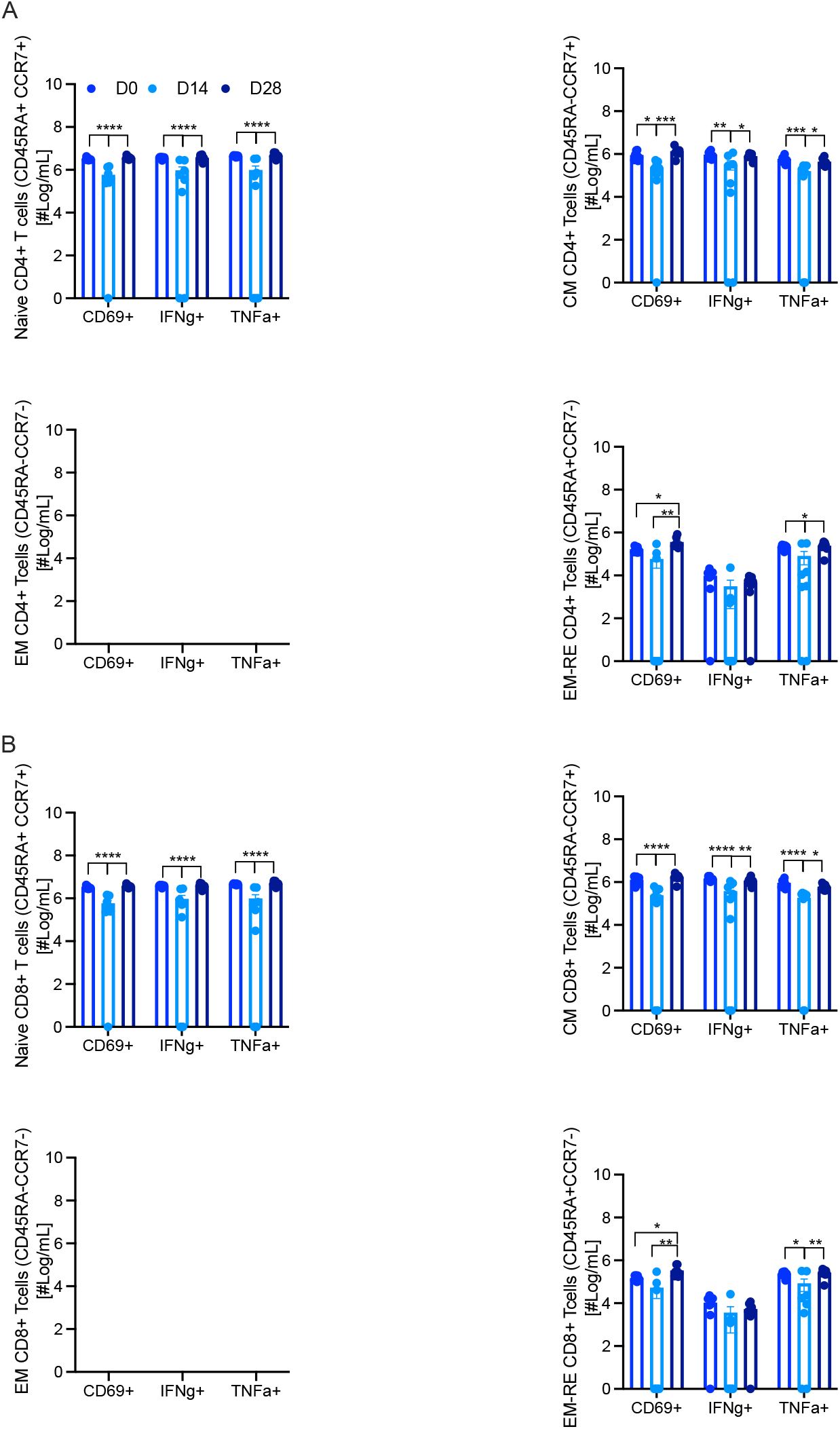
T cell responses after MARV infection in NHPs vaccinated 7 days before challenge. Number of circulating MARV GP-specific activated T cells. (**A**) CD4^+^ or (**B**) CD8^+^ T cells representing naïve, central memory (CM), effector memory (EM), and effector memory re-expressing (EM-RE)populations expressing CD69, IFNγ, and TNFα. Mean and SEM are depicted. Statistical significance as determined by the Mann–Whitney test is indicated as *p* < 0.0001 (****), *p* < 0.001 (***), *p* < 0.01 (**), and *p* < 0.05 (*).

**Figure S4.**
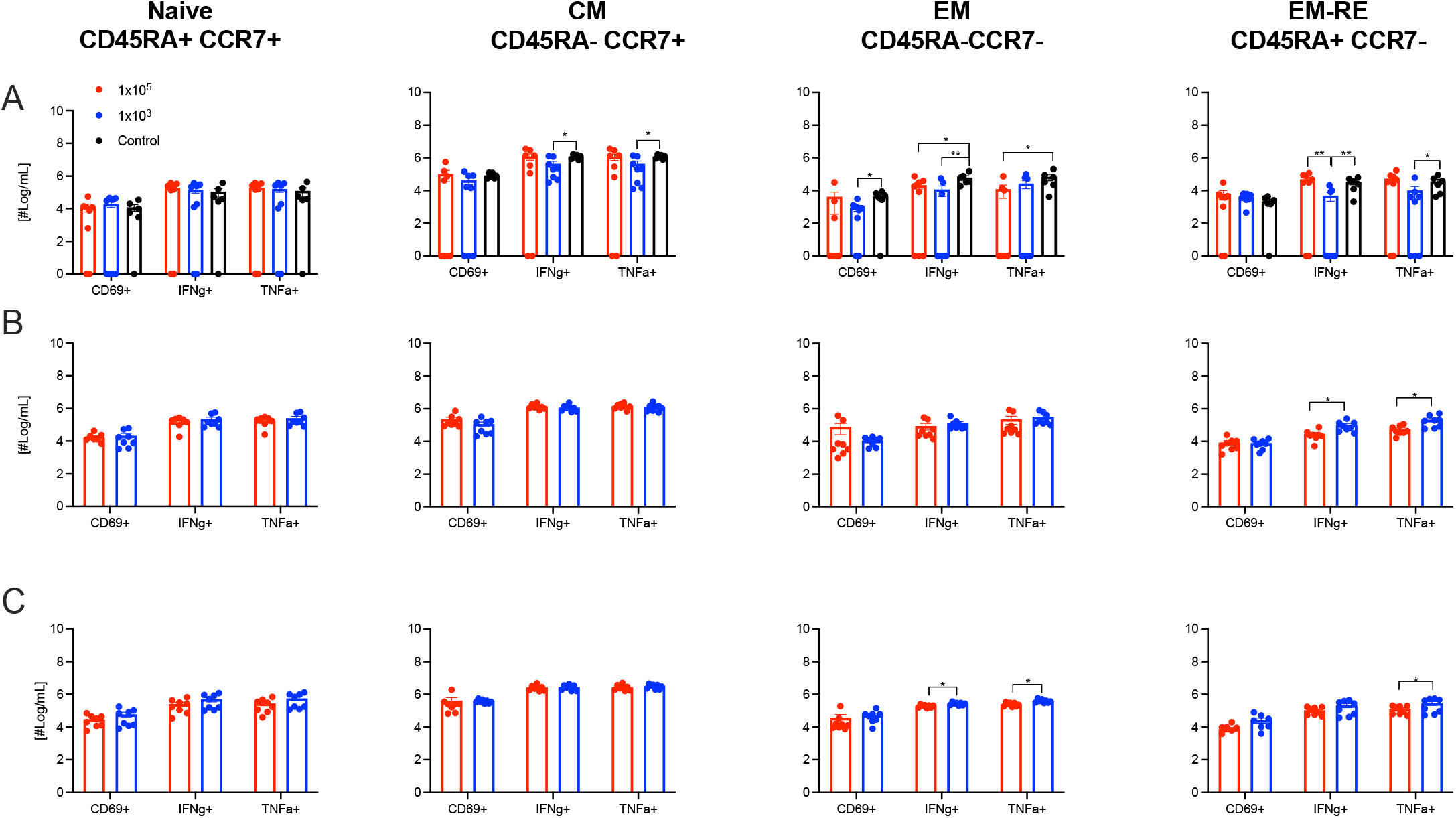
CD8^+^ T cell responses after MARV infection in NHPs vaccinated 14 days before challenge. Number of circulating MARV GP-specific activated T cells on (**A**) 0 DPC, (**B**) 14 DPC, and (**C**) 28 DPC. Populations represent naïve, central memory (CM), effector memory (EM), and effector memory re-expressing (EM-RE) cells expressing CD69, IFNγ, and TNFα. Mean and SEM are depicted Mean and SEM are depicted. Statistical significance as determined by the Mann–Whitney test is indicated as *p* < 0.01 (**), and *p* < 0.05 (*).

## Notes

### Competing Interest Statement

The authors have declared no competing interest.

